# scGAE: topology-preserving dimensionality reduction for single-cell RNA-seq data using graph autoencoder

**DOI:** 10.1101/2021.02.16.431357

**Authors:** Zixiang Luo, Chenyu Xu, Zhen Zhang, Wenfei Jin

## Abstract

Dimensionality reduction is crucial for the visualization and interpretation of the high-dimensional single-cell RNA sequencing (scRNA-seq) data. However, preserving topological structure among cells to low dimensional space remains a challenge. Here, we present the single-cell graph autoencoder (scGAE), a dimensionality reduction method that preserves topological structure in scRNA-seq data. scGAE builds a cell graph and uses a multitask-oriented graph autoencoder to preserve topological structure information and feature information in scRNA-seq data simultaneously. We further extended scGAE for scRNA-seq data visualization, clustering, and trajectory inference. Analyses of simulated data showed that scGAE accurately reconstructs developmental trajectory and separates discrete cell clusters under different scenarios, outperforming recently developed deep learning methods. Furthermore, implementation of scGAE on empirical data showed scGAE provided novel insights into cell developmental lineages and preserved inter-cluster distances.

## Introduction

Single-cell RNA sequencing (scRNA-seq) is an ideal approach for investigating cell-cell variation. Conventional dimensionality reduction techniques such as principal component analysis (PCA) and t-Distributed Stochastic Neighbor Embedding (t-SNE)^1^ were implemented on scRNA-seq data for visualization and downstream analyses, significantly increasing our understanding of cellular heterogeneity and development progress. The recent emergence of massively parallel scRNA-seq such as droplet platforms enabled interrogation of millions of cells in complex biological systems^2–5^, which provide a fantastic potential for dissection of tissue and cellular microenvironment, identification of rare/new cell types, inference of developmental lineages, and elucidation of the mechanism of cellular response to stimulations^6^. However, the data generated by massively parallel scRNA-seq are of high dropout and high noise with complex structure, which posed a series of challenges on dimensionality reduction. Particularly, it is a big challenge to preserve the complex topological structure among cells.

Many dimensionality reduction methods have been developed or introduced for scRNA-seq data analyses in the past several years. Recently developed competitive methods include DCA^7^, SCVI^8^, scDeepCluster^9^, PHATE^10^, SAUCIE^11^, and Ivis^12^. Among them, deep learning showed the greatest potentials. For instance, DCA, scDeepCluster, Ivis, and SAUCIE adapted the autoencoder to denoise, visualize and cluster the scRNA-seq data. However, these deep learning-based models only embedded the distinct cell features while ignoring the cell-cell relationships, which limited their ability to reveal the complex topological structure among cells and made them difficult to elucidate the developmental trajectory. The recently proposed graph autoencoder^13^ is very promising as it preserves the long-distance relationships among data in a latent space. In this study, we developed the single-cell graph autoencoder (scGAE). It improved the graph autoencoder to preserving global topological structure among cells. We further extended the scGAE for visualization, trajectory inference, and clustering. Analyses of simulated data and empirical data showed that scGAE outperformed the other competitive methods.

## Results

### The Model architecture of scGAE

scGAE combines the advantage of the deep autoencoder and graphical model to embed the topological structure of high-dimensional scRNA-seq data to a low-dimensional space (Fig1). After getting the normalized count matrix, scGAE builds the adjacency matrix among cells by K-nearest-neighbor algorithm. The encoder maps the count matrix to a low-dimensional latent space by graph attentional layers^14^. scGAE decodes the embedded data with a feature decoder and a graph decoder. The feature decoder reconstructs the count matrix to preserve the feature information; The graph decoder recovers the adjacency matrix and preserves the topological structure information. It decodes the embedded data to the spaces with the same dimension as original data by minimizing the distance between the input data and the reconstructed data (see Methods). We use deep clustering to learn the data embedding and do cluster assignment simultaneously^15^, generating a clustering-friendly latent representation. The implementation and usage of scGAE can be found on Github: https://github.com/ZixiangLuo1161/scGAE.

**Figure 1.**
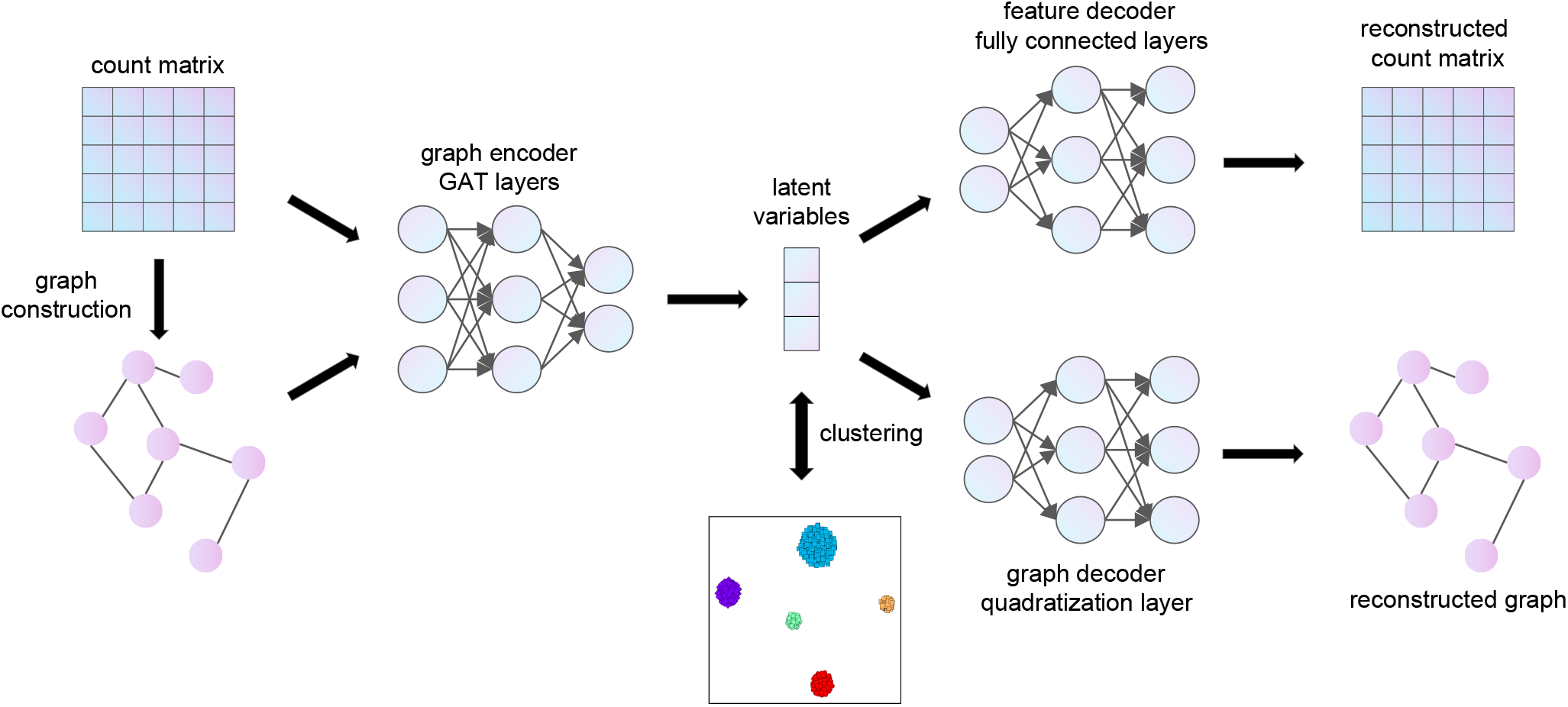
The Model architecture of scGAE. The normalized count matrix represents the gene expression level in each cell. The adjacency matrix is constructed by connecting each cell to its K nearest neighbors. The encoder takes the count matrix and the adjacency matrix as inputs and generates low-dimensional latent variables. The feature decoder reconstructs the count matrix. The graph decoder reconstructs the adjacency matrix. Clustering is performed on the latent variables.

### Visualization of scGAE embedded data and comparison to other methods

To systematically evaluate the performance of scGAE, we summarized four representative scenarios (scenario1: cells in continuous differentiation lineages; scenario2: cells in differentiation lineages where cells concentrate at the center of each branch; scenario3: distinct cell populations with apparent differences; and scenario4: distinct cell populations with small population differences) (Fig2 left). We used Splatter^16^ and PROSSTT^17^ to simulate scRNA-seq data for scenario1, scenario2, scenario3, and scenario4. The latent embedding inferred by scGAE was visualized by tSNE. In scenario1 and secnario2, scGAE almost entirely reproduced the differentiation lineages (Fig2a, 2b), while other methods only revealed some local structures and failed to exhibit the overall structure of simulated data (Fig2a, 2b). The results of tSNE and SAUCIE exhibited distinct clusters but lost lineage relationship in scenario2 (Fig2b). In scenario3 and secnario4, scGAE almost perfectly preserved the compact cell clusters and inter-cluster distances in the simulated data, while the clusters inferred by other methods are dispersed, and the topological structure among these clusters was not preserved (Fig2c, 2d, Supplemental figure 1). Only scGAE separated all the clusters while the other methods mixed different types of cells when the differences between clusters are small (Fig 2d). Based on these observations, scGAE perfectly reproduced the differentiation lineages and distinct clusters in the simulated data (Fig2), indicating scGAE outperforms other competitive methods in restoring the relationship between cells.

**Figure 2.**
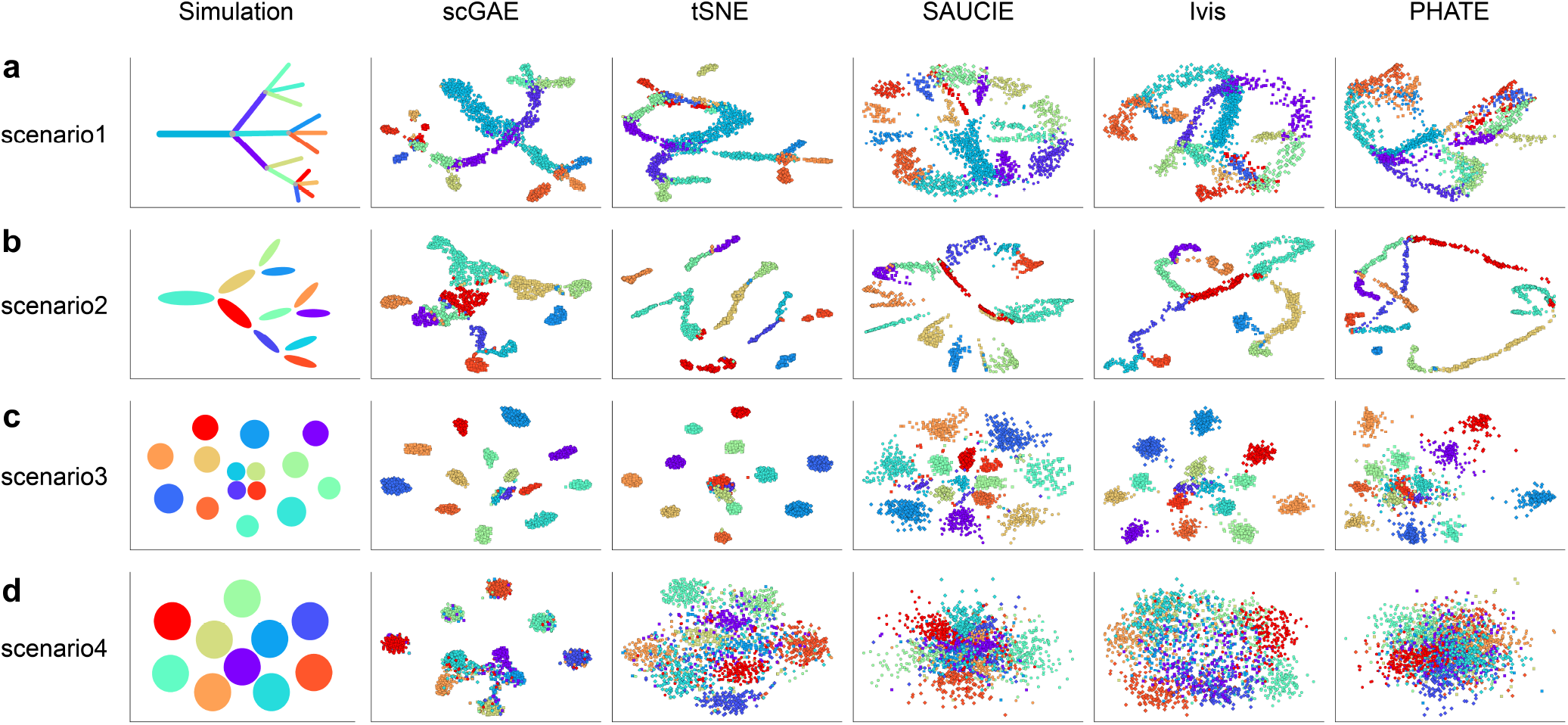
Visualization of the four simulated datasets by scGAE, tSNE, SAUCIE, Ivis, and PHATE. Each color represents a cell subpopulation in the simulated dataset. (a) scenario1: cells in continuous differentiation lineages. (b) scenario2: cells in differentiation lineages where cells concentrate at the center of each branch. (c) scenario3: distinct cell populations with apparent population differences. (d)scenario4: distinct cell populations with small population differences.

### Trajectory inference and cell clustering based on scGAE embedded data

We further quantitatively evaluated the performance of scGAE for trajectory inference tasks. The scGAE and several other competitive methods were used to perform dimensionality reduction on the simulated lineages (simulated by PROSSTT) (scenario1 and 2). We conducted trajectory inference on these embedded data using DPT^18^. The Kendall correlation coefficient^19^ between the inferred trajectories and the ground truth was calculated to measure their similarity. The results showed that scGAE and SCVI better recovered the original trajectory than the other competitive methods on both scenario1 and 2 (Fig3a, Fig3b). Next, we evaluated the performance of scGAE on cell clustering tasks. Simulated data with cell clusters (simulated by Splatter) (scenario3 and 4) were analyzed by scGAE and other competitive methods. We performed Louvain clustering on these embedded data. Normalized mutual information (NMI) was used to measure the difference between inferred clusters and ground truth. The results showed that scGAE was the best among these methods (Fig3c, Fig3d). Although SCVI is the second-best performed for trajectory inference (Fig3a, Fig3b), it is the worst performed for cell clustering (Fig3c, Fig3d). On the other hand, PCA is the second-best method for cell clustering (Fig3c, Fig3d), while it does not perform well for trajectory inference. Overall, scGAE performed best for both trajectory inference and cell clustering.

**Figure 3.**
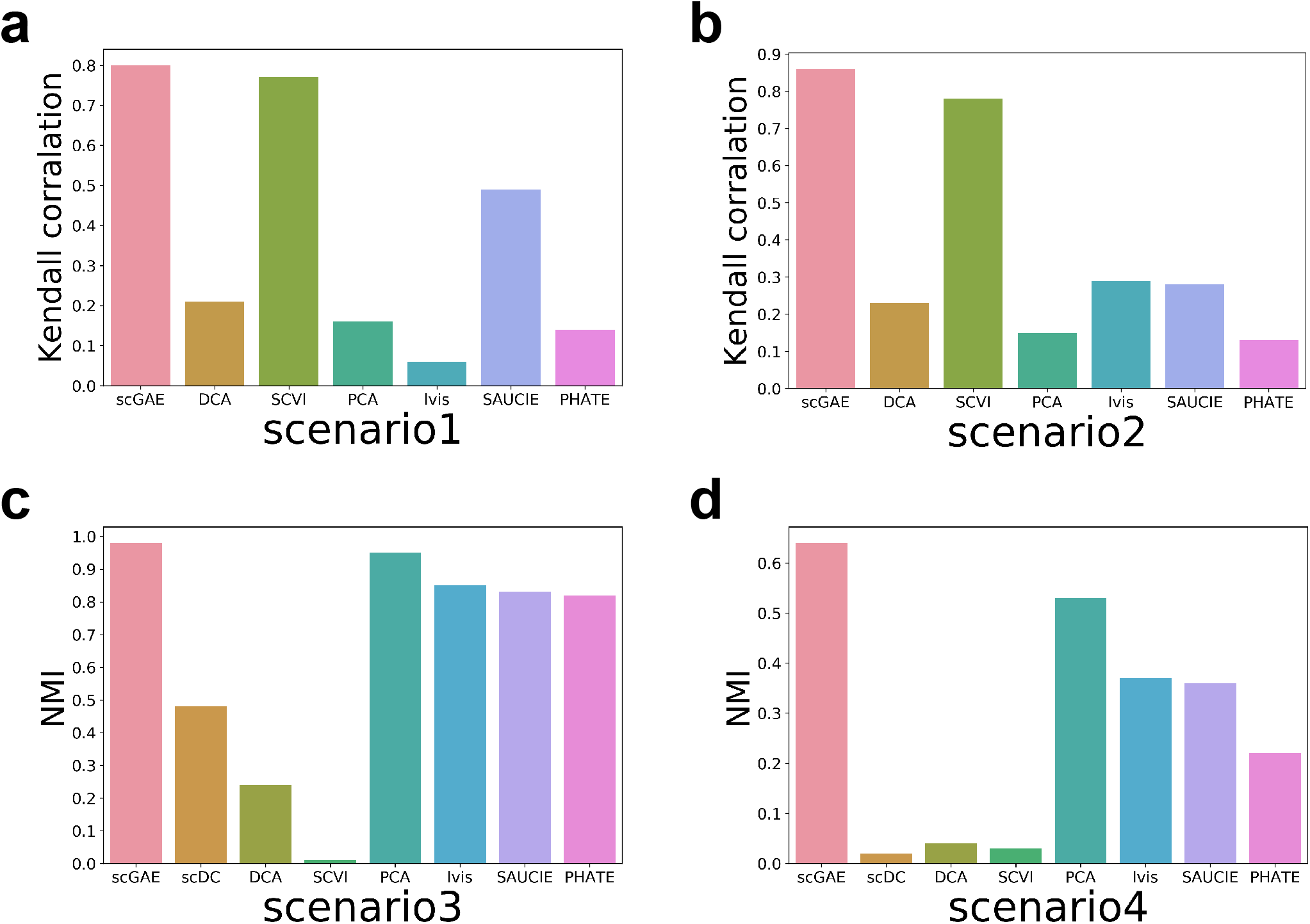
Quantitative evaluation of scGAE and several other competitive methods on clustering and trajectory inference tasks. In scenario1 (a) and scenario2 (b), the Kendall correlation between the ground truth and inferred trajectory was calculated. In scenario3 (c) and scenario4 (d), the normalized mutual information (NMI) measures the difference between the ground truth and the inferred clusters.

### scGAE identified novel subpopulations that shaped hematopoietic lineage relationship

Single cell analysis of hematopoietic stem and progenitor cells (HSPCs) have significantly increased our understanding of the early cell subpopulations and developmental trajectory during hematopoiesis^5, 20–25^.We further used scGAE to analyze HSPCs scRNA-seq data from our previous study^5^ (Fig4a). We found the previous identified Basophil/Eosinophil/Mast progenitors (Ba/Eo/MaP) has been classified into multiple subpopulations (Fig4b). It indicates that the cells in Ba/Eo/MaP may have different differentiation potentials at early phase. While the other competitive methods did not identify the subpopulations in Ba/Eo/MaP (Supplemental figure 2), supporting scGAE has the highest statistical power to identify the substructure in the scRNA-seq data.

**Figure 4.**
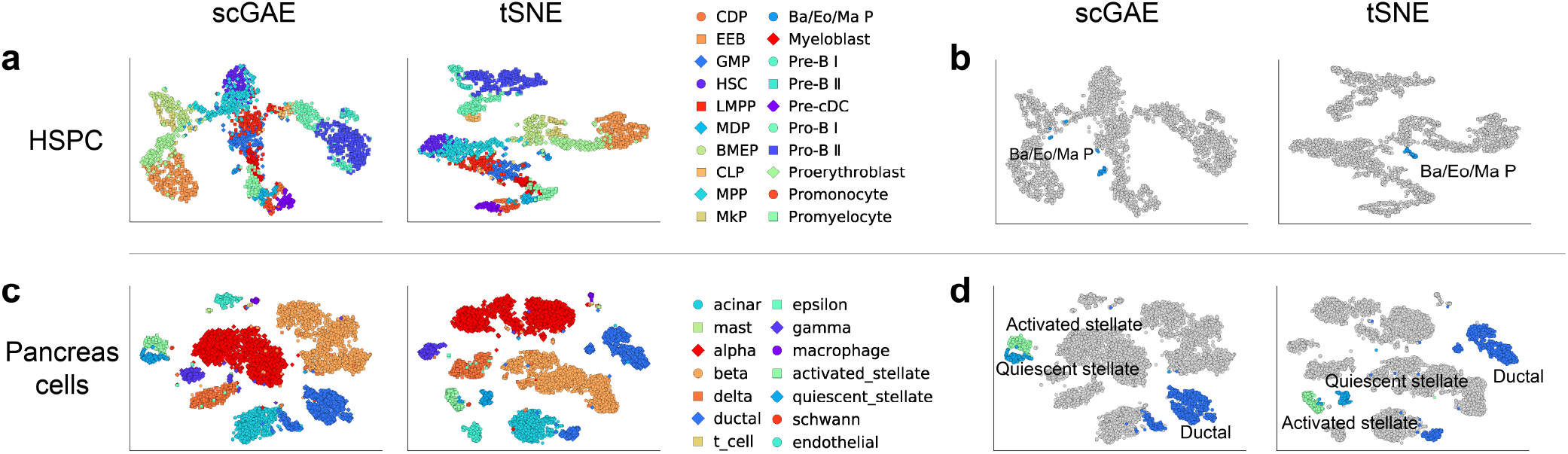
Analyses of two real datasets. (a)Visualization of HSPC cells by scGAE and tSNE (b) scGAE identified the multiple subpopulations in previous reported Ba/Eo/MaP. (c) Visualization of pancreases cells by scGAE and tSNE. (d) The close distance between two stellate states and the short distance between ductal subtypes recovered by scGAE.

### scGAE preserved topological structure among human pancreatic cells populations

The function of the pancreas hinges on complex interactions among distinct cell types and cell populations. We re-analyzed the scRNA-seq data of human pancreatic cells from Baron et al.^26^. Although the pancreatic cell subpopulations identified by scGAE are the same as the original study, we found the distances and topological structures among cell types inferred by scGAE better fit our knowledge (Fig4c). For instance, the activated stellate and quiescent stellate showed similar expression profiles and are very close to each other^27^, which is recovered by scGAE while not recovered by other methods (Fig4d and Supplemental figure 2). scGAE also preserved the short distance between two ductal subtypes, while other methods did not (Fig4d and Supplemental figure 2). Overall, scGAE preserved the topological structure among different cell populations, which greatly benefit our understanding of the cellular relationships.

## Discussion

Because of the high noises of scRNA-seq data and complicated cellular relationships, preserving the topological structure of scRNA-seq data in low-dimensional space is still a challenge. We proposed scGAE which is a promising topology-preserving dimensionality reduction method. It generates a low-dimensional representation that better preserves both the global structure and local structure of the high-dimensional scRNA-seq data. The key innovation of scGAE is to embed the structure information and feature information simultaneously using a multitask graph autoencoder. It is suitable for analyzing the data both in lineages and clusters. The learned latent representation benefits various downstream analyses, including clustering, trajectory inference, and visualization. The analyses on both simulated data and empirical data suggested scGAE accurately preserved the topological structures of data.

As the first study adapting graph autoencoder for dimensionality reduction of scRNA-seq data, this approach is likely to be significantly improved in the future. Firstly, because the complex data structure is hard to be directly embedded into two-dimensional space by graph autoencoder, we embedded the scRNA-seq data into an intermediate dimension and used tSNE to visualize the embedded data into a two-dimensional space. However, the tSNE focuses more on local information, and it sometimes fails to correctly recover the global structure, which may distort the topological structure in the data. A better visualization method is needed to preserve the topological structure of scRNA-seq data. Secondly, the graph in scGAE is constructed by the K-nearest neighbor (KNN) algorithm that relies on a predefined parameter K. However, the optimal K varies among different datasets and different parts of a dataset. Constructing an optimal graph is challenging due to the difficulty in determining a suitable K, which could be our potential future endeavors.

## Methods

### Joint graph autoencoder

The graph autoencoder is a type of artificial neural network for unsupervised representation learning on graph-structured data^13^. The graph autoencoder often has a low-dimensional bottleneck layer so that it can be used as a model for dimensionality reduction. Let the inputs be single-cell graphs of node matrices *X* and adjacency matrices *A*. In our joint graph autoencoders^28^, there is one encoder *E* for the whole graph and two decoders *D*_*X*_ and *D*_*A*_ for nodes and edges respectively. In practice, we first encode the input graph into a latent variable *h* = *E*(*X, A*), and then we decode *h* into the reconstructed node matrix *X*_*r*_ = *D*_*X*_ (*h*) and the reconstructed adjacency matrix *A*_*r*_ = *D*_*A*_(*h*). The objective of learning process is to minimize the the reconstruction loss

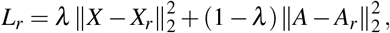

where the weight *λ* is a hyper-parameter. In our experiments, *λ* is set to be 0.6.

We used the Python package Spektral^29^ to implement our model. There are many types of graph neural networks that can be used as the encoder or decoder. Hereby, to extract the features of a node with the aid of its neighbors, we apply graph attention layers as default in the encoder. Other graph neural networks such as GCN^30^, GraphSAGE^31^ and TAGCN^32^ can also be implemented as the encoder in scGAE. The feature decoder *D*_*X*_ is a four-layer fully connected neural network with 64, 256, 512 nodes in hidden layers.

The edge decoder consists of a fully connected layer followed by the composition of quadratization and activation:

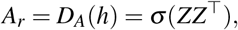

where *Z* = *σ* (*Wh*) arises as an output of a fully connected layer with the weight matrix *W*, and *σ* (*x*) = max(0*, x*) is the rectified linear unit.

### Deep-clustering embedding

Motivated by Yang et al^33^, we use a two-stage method. The first stage is to pre-train scGAE by minimizing *L*_*r*_. The resulting neural network parameters are set as the initialization of the second stage, which we call alter-training. The loss function in the alter-training stage compromises both reconstruction error *L*_*r*_ and clustering cost *L*_*c*_ = *L*_*c*_(*h, μ*):

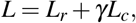

where *μ* is a collection of clustering centroids, and *γ* is a hyper-parameter set as 2.5 in our experiments.

The alter-training consists of doing the following two steps alternately:

1. Given a collection of clustering centroids *μ*, update network parameters by minimizing *L*;
2. Compute the embedded data *h* using the updated network, and do clustering in the embedded space to obtain new centroids *μ*;

In experiments, we use the pre-trained network to generate the initial embedded data which are clustered to obtain the initial centroids by Louvain^34^. There are various choices for the loss *L*_*c*_ and the clustering algorithm in the second step^15^. In practice, we compute the new centroids *μ* by minimizing *L*_*c*_ using the stochastic gradient descent. A good choice of *L*_*c*_ is the soft assignment loss^35^, which is the KL divergence of empirical clustering assignment distribution *Q* from a target distribution *P*.

Given an embedded point *h*_*i*_ and a centroid *μ*_*j*_, *Q* is defined as Student’s t-distribution 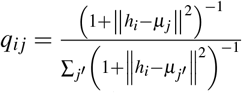. An ideal target distribution should have the following properties: (1) improve cluster purity, (2) put more emphasis on data points assigned with high confidence, and (3) prevent large clusters from distorting the hidden feature space. In experiments, we choose *P* as 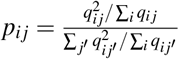.

### Evaluation metric

Clustering results are measured by Normalized Mutual Information (NMI)^36^. Given the knowledge of the ground truth class assignments *U* and our clustering algorithm assignment *V* on *n* data points, NMI measures the agreement of the two assignment, ignoring permutations. NMI is defined as

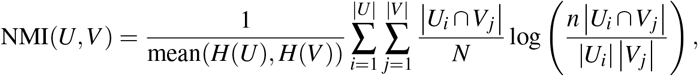

where 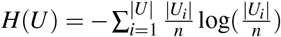 is the entropy.

Trajectory inference results are measured by Kendall correlation coefficient. We define an order among the set of observations (*x*_1_*, y*_1_), (*x*_2_*, y*_2_)*, …,* (*x*_*n*_, *y*_*n*_): any pair of observations (*x*_*i*_, *y*_*i*_) and (*x*_*j*_, *y*_*j*_), where *i < j* are said to be concordant if either both *x*_*i*_ > *x*_*j*_ and *y*_*i*_ > *y*_*j*_ hold or both *x*_*i*_ < *x*_*j*_ and *y*_*i*_ < *y*_*j*_ hold; otherwise they are said to be discordant. Denote the number of concordant pairs as *N*_*conco*_ and the number of discordant pairs as *N*_*discon*_, Kendall correlation coefficient is defined as

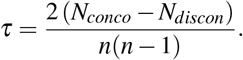

### Data simulation

We simulated five scRNA-seq datasets using Splatter R package (data1, data3, and data4) and PROSSTT Python package (data2 and data5). The cells in data1 and data5 are in the linear distribution along the developmental trajectory. The cells in data2 have a skewed distribution where cells concentrate at the center of each branch. The cells in data3 and data4 are in distinct clusters with moderate and small cluster differences, respectively. All datasets have 2000 cells and 5000 genes. Data1, data2, data3, and data4 were simulated for scenario1 to scenario4 for data visualization. Data5, data2, data3, and data4 are used for the evaluation of scGAE on trajectory inference and cell clustering tasks.

### Data preprocessing

The scRNA-seq data preprocessing was conducted using scTransform^37^ in The Seurat package^38^. The pre-processed count matrix was used to construct the single-cell graph, where the nodes represent cells, and the edges represent the relationships between cells. The cell graph is built by the K-nearest neighbor (KNN) algorithm^39^ in the Scikit-learn Python package^40^. The default K is predefined as 35 in this study and adjusted according to the datasets in our experiments. The generated adjacency matrix is a 0-1 matrix, where 1 represents being connected, and 0 represents no connection.

### Empirical scRNA-seq data

We analyzed two different scRNA-seq datasets, namely HSPCs data and pancreatic cells data. HSPCs data and pancreatic cells data represent cells showing lineages relationship and cells showing distinct clusters, respectively. The HSPCs data are single-cell transcriptome data of FACS sorted CD34+ cells from human bone marrow mononuclear cells, accessible in the national genomics data center (HRA000084) and described in our previous study^5^. The pancreases cells data contains 10,000 single-cell transcriptomes with 14 distinct cell clusters, download from GEO (GSE84133)^26^.

### Competitive methods

Seven competitive methods, namely scDeepCluster, DCA, SCVI, PCA, Ivis, SAUCIE, and PHATE, were compared with scGAE. Among these methods, scDeepCluster, DCA, SCVI, Ivis, and SAUCIE are deep learning based and showed the greatest potential. These methods usually generate hidden variables for downstream analysis, including visualization, clustering, and trajectory inference. The raw count matrix was used as input for DCA, SCVI, and scDeepCluster. For methods that take normalized data as input (scGAE, SAUCIE, PCA, Ivis, and PHATE), scTransform was used for data preprocessing. Each software was run following its manual and with default parameters. For DCA, PCA was conducted to reduce the DCA-denoised data to 32 PCs. For SAUCIE and Ivis, PCA reduced the preprocessed data to 100 PCs and 50 PCs, respectively. Ivis, SAUCIE, and PHATE directly generate the 2-dimensional embeddings. The cell clustering and trajectory inference were performed on the two-dimensional embeddings. Both scGAE and PCA embedded simulated data to 10 dimensions and embedded empirical data to 20 dimensions due to the complex structure of the empirical data.

## Acknowledgements

This study was supported by National Key R&D Program of China (2018YFC1004500), National Natural Science Foundation of China (81872330, 31741077), the Science and Technology Innovation Commission of the Shenzhen Municipal Government (JCYJ20170817111841427), the Shenzhen Science and Technology Program (KQTD20180411143432337), Center for Computational Science and Engineering of Southern University of Science and Technology.

## Author contributions statement

W.J. and Z.Z. conceived and designed the project. Z.L. and C.X. developed the algorithm, coded the program and performed the data analysis. W.J. and Z.L. wrote the manuscript with inputs from all authors.

## Additional information

### Accession codes

The code and software of scGAE are available on GitHub (https://github.com/ZixiangLuo1161/scGAE);

